# Gasotransmitter H_2_S accelerates seed germination via activating AOX mediated cyanide-resistant respiration pathway

**DOI:** 10.1101/2021.10.13.464324

**Authors:** Huihui Fang, Ruihan Liu, Zhenyuan Yu, Gang Wu, Yanxi Pei

**Affiliations:** Laboratory of Plant Molecular and Developmental Biology, Collaborative Innovation Center for Efficient and Green Production of Agriculture in Mountainous Areas of Zhejiang Province, College of Horticulture Science, Zhejiang Agriculture and Forestry University, Hangzhou, Zhejiang, 311300, China; School of Life Science and Shanxi Key Laboratory for Research and Development of Regional Plants, Shanxi University, Taiyuan, Shanxi, 030006, China

**Keywords:** Gasotransmitter, Hydrogen sulfide, Alternative oxidase, Cyanide-resistant pathway, Cell reducing power, Seed germination

## Abstract

Hydrogen sulfide (H_2_S) has been witnessed as a crucial gasotransmitter involving in various physiological processes in plants. H_2_S signaling has been reported to involve in regulating seed germination, but the underlying mechanism remains poorly understood. Here, we found that endogenous H_2_S production was activated in germinating Arabidopsis seeds, correlating with upregulated both the transcription and the activity of enzymes (LCD and DES1) responsible for H_2_S production. Moreover, NaHS (the H_2_S donor) fumigation significantly accelerated seed germination, while H_2_S-generation defective (*lcd/des1*) seeds exhibited decreased germination speed. Further results indicated that the alternative oxidase (AOX), a cyanide-insensitive terminal oxidase, can be stimulated by imbibition, and the expression of *AOX* genes was provoked lag behind H_2_S production during germination. Additionally, exogenous H_2_S fumigation significantly reinforced imbibition induced enhancement of *AOX1A* expression, and mediated post-translational modification to keep AOX in its reduced and active state, which mainly involved H_2_S induced increase of the GSH/GSSG ratio and the cell reducing power. Consequently, H_2_S signaling acts as a trigger to induce AOX mediated cyanide-resistant respiration to accelerate seed germination. Our study correlates H_2_S signaling to cyanide metabolism, which also participates in endogenous H_2_S generation, providing evidence for more extensive studies of H_2_S signaling.

**Highlight:** Gasotransmitter H_2_S provokes AOX mediated cyanide-resistant respiration, mainly through both long-term (up-regulating *AOX1A* expression) and short-term (inducing post-translational activation of AOX) regulatory modes, to accelerate seed germination.

## 1. Introduction

Seed germination, a crucial stage of plant life cycle, is of major importance for early seedling emergence and is the first step to achieve high crop yield and quality (Rajjou *et al*., 2012; Finch-Savage and Bassel, 2016), so understanding the potential molecular mechanism of seed germination has great significance. Hydrogen sulfide (H_2_S), alongside nitric oxide (NO) and carbon monoxide (CO), has been recognized as a multifunctional gasotransmitter that plays crucial roles in plants development and stresses responses (Wang, 2002; Yang *et al*., 2004; Arif *et al*., 2021; Liu *et al*., 2021; Zhang *et al*., 2021). The possible involvement of H_2_S signaling in seed germination has been reported (Baudouin *et al*., 2016; Zhou *et al*., 2018; Chen *et al*., 2019); however, the underlying metabolism has not been thoroughly elucidated to date.

In plants, both nonenzymatic and enzymatic pathways are responsible for H_2_S generation, although the nonenzymatic pathway only accounts for a small portion of H_2_S sources (Jin and Pei, 2015). Enzymes that produce endogenous H_2_S in plants can be roughly divided into two major categories, the cysteine desulfhydrases (CDes) and O-acetyl-L-serine (thiol) lyase (OASTL). For the former case, CDes degrade cysteine into H_2_S, ammonia, and pyruvate in a stoichiometric ratio of 1:1:1, while for the latter case, free H_2_S appears to be released only in a side reaction of the incorporation of inorganic S into cysteine mediated by OASTL (Liu *et al*., 2021; Zhang *et al*., 2021). Specially, Alvarez *et al*. (2010) characterized a novel L-cysteine desulfhydrase (EC 4.4.1.1) DES1, which is an O-acetylserine(thiol)lyase homolog based on its sequence feature but exhibits higher CDes activity and has a much higher affinity to L-cysteine, mediating the generation of H_2_S in the cytosol (Alvarez *et al*., 2010). Even more to the point, the β-cyanoalanine synthase (β-CAS), which mediates a principal route for cyanide metabolism in plants, could also induce H_2_S generation by catalyzing the conversion of cysteine and cyanide to H_2_S and β-cyanoalanine, a process that acting as an important linker in regulating cyanide detoxification and H_2_S generation (Garcia *et al*., 2010; Romero *et al*., 2014).

The potential physiological and molecular role of cyanide in plants development and stress responses has been noted, although most cyanide would be rapidly detoxified and metabolized to keep its concentration below toxic levels (Garcia *et al*., 2010). Several pieces of research have suggested the emission of cyanide during the pre-germination period of many seeds (Bethke *et al*., 2006; Esashi *et al*., 2006), and the released cyanide in germinating seeds may help break dormancy and promote germination (Oracz *et al*., 2009; Gniazdowska *et al*., 2010; Oracz *et al*., 2008; Dobrzynska *et al*., 2005). It is well known that cyanide binds irreversibly to the heme iron of terminal cytochrome c oxidase (COX) in the mitochondrial electron transport chain, thereby blocking electron transfer from reduced cytochrome c to oxygen (Solomonson, 1981; Vennesland, 1981). However, an alternative electron-transfer pathway is insensitive to cyanide, which is known as cyanide-resistant respiration pathway. Correspondingly, the cyanide-resistant respiration, a respiration alternative pathway, has been reported to be triggered by seed imbibition (Esashi *et al*., 1979; Burguillo and Nicolas, 1977). Subsequently, a pending question of the relationship between H_2_S signaling and cyanide-resistant respiration during seed germination attracts our attention.

Cyanide-resistant respiration involves a cyanide-insensitive terminal oxidase, the alternative oxidase (AOX) (LATIES and GG, 1982; Vanlerberghe *et al*., 1994), which provides a parallel pathway for mitochondrial electron flow, bypassing complexes III and IV, and results in cyanide-resistant respiration. To a certain extent, the AOX activity could represent the degree of the involvement of cyanide-resistant respiration. AOX is a nuclear-encoded mitochondrial protein, which exists as a dimer in the inner mitochondrial membrane and has two states, the oxidized state in which the dimer is covalently cross-linked by a disulfide bridge (-S-S-), and the reduced state (-SH HS-) maintained through non-covalent interaction. The reduced AOX form can be four- to five-fold more active than the oxidized dimeric AOX (Moore *et al*., 1995; Sluse and Jarmuszkiewicz, 1998; Day and Wiskich, 1995), so the activity of AOX can be regulated by its redox state and the ratio of reduced form to oxidized dimeric form.

In this study, the correlation between H_2_S signaling and AOX mediated cyanide-resistant respiration during seed germination was widely explored. We found that H_2_S could enhance the cyanide-resistant respiration in germinating seeds by reinforcing the increase of *AOX1A* expression and mediating post-transcriptional modification on AOX to maintain more AOX in its reduced and active state, which provides molecular evidence for understanding the mechanism of the gasotransmitter H_2_S in the acceleration of seed germination by activating the cyanide-resistant respiration. Our study provides a body of evidence for demonstrating the function of H_2_S signaling in regulating respiratory pattern in germinating seeds, and correlates the H_2_S signaling to cyanide metabolism, which also participates in endogenous H_2_S generation, providing evidence for more extensive studies of H_2_S signaling.

## 2. Materials and Methods

### 2.1 Plant Materials and treatments

The seeds of *A. thaliana* Wild type (WT, Columbia, Col-0), the *lcd/des1* double mutants (*lcd/des1*), and the transgenic plants over-expressing DES1 (OE-*DES1*) were used in this study. The *lcd/des1* double mutant was obtained by crossing the *LCD* T-DNA insertion mutant *lcd* (SALK_082099) with the *DES1* defective mutant *des1* (SALK_205358C). The DES1 over-expression transgenic plants (OE-*DES1*) were generated as described previously (Zengjie *et al*., 2015).

For germination tests, seeds were stratified at 4°C for 48 h, then germinated in the Petri dishes on 3 layers of filter papers soaked by ½ Murashige-Skoog (½ MS) fluid medium, 100 seeds per dish, and each kind of seeds or treatment had 3 replicates. All dishes were subsequently kept in a 16 h/8 h (light/dark) photoperiod with a light illumination of 160 Em^-2^s^-1^ at 23°C and 60% relative humidity. The seed germination was analyzed over time, and the germination percentage was assessed by measuring the rate of testa ruptured seeds, described in the Fig. 5 (Fig. 5).

For exogenous H_2_S fumigation treatment, the seeds were successively fumigated with H_2_S released by the donor NaHS immediately after being placed on the filter paper. The NaHS solution-containing tube was placed into the Petri dishes, which were then sealed with parafilm. Every fumigation lasted for 12 hours, with an interval of 24 hours between fumigations.

### 2.2. Measurement of the H_2_S production rate and endogenous H_2_S content

To confirm whether the endogenous H_2_S signaling participates in regulating seed germination, the endogenous H_2_S production rate and H_2_S content in germinating seeds were measured. Total protein extracts were collected from seeds at various time points after water imbibition, then cysteine was added as a substrate in the extracts to determine the production rate of H_2_S mediated by CDes according to previously described methods (Fang *et al*., 2017; Jin *et al*., 2013). The endogenous H_2_S content was detected using a novel polarographic H_2_S sensor (WPI, TBR4100, Sarasota, FL, USA). Briefly, 0.1 g seeds were frozen and ground in liquid nitrogen, then 1 mL PBS buffer (0.05 mol L^-1^, pH 6.8, containing 0.2 mol L^-1^ AsA and 0.1 mol L^-1^ EDTA), was added to the well-ground powder, then the suspensions were taken to determine H_2_S content according to a previous publication (Liu *et al*., 2019).

### 2.3 Total RNA extraction and real-time quantitative RT-PCR

Total RNA was extracted from imbibed seeds using RNAiso plus reagent (TaKaRa, Shiga, Japan, Cat9109) according to the manufacturer’s instructions. The cDNA was synthesized using a reverse transcription system kit (PrimeScript RT Reagent Kit, TaKaRa, RR037B) and oligo(dT) primers, and then the real-time quantitative RT-PCR (qRT-PCR) was performed to detect the mRNA level of target genes according to the instructions of the Bio-Rad Real-Time System (CFX96TM C1000 Thermal Cycler), then the relative expression of target genes was calculated by using the 2^−δδCt^ method. All of the primer pairs used for qRT-PCR were checked for amplification specificity and were listed in Table 1. Ubiquitin4 (UBQ4, At5g20620) was used as the internal control.

**Table 1.**
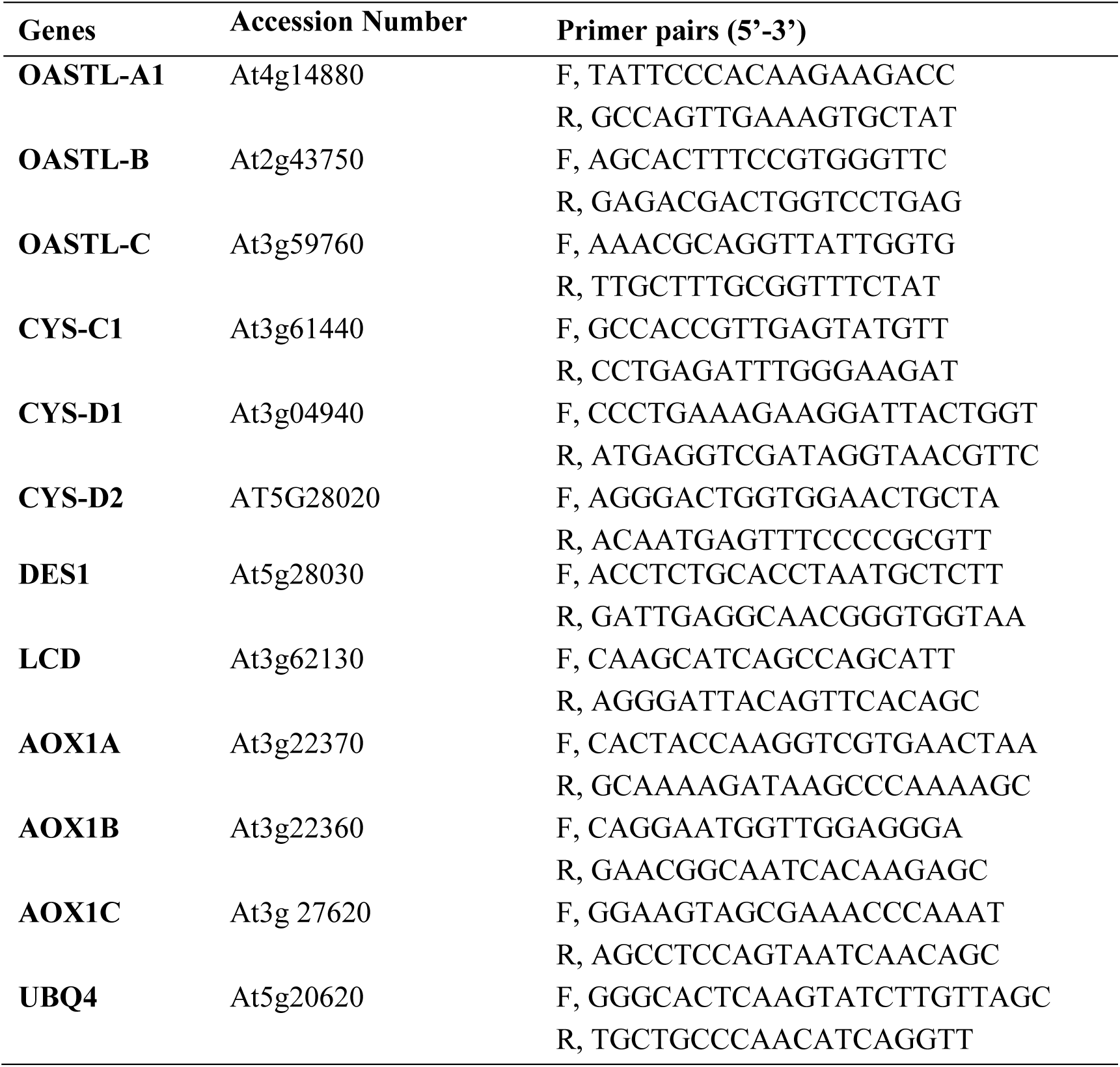
List of all primers for qRT-PCR used in this study.

### 2.4 Determination of the GSH and GSSG content

The contents of the reduced GSH and oxidized form GSSG were measured using a GSH and GSSG assay kit S0053 (Beyotime Institute of Biotechnology, China) based on a previously described method (Fang *et al*., 2014), and then the ratio of GSH/GSSG was calculated.

### 2.5 Assay of the AOX concentration and its redox state

The AOX concentration during seed germination was determined using the plant alternative oxidase (AOX) ELISA Kit (YS07118B, GTX, China) according to the manufacturer’s instructions. The AOX concentration was then calculated by comparing the optical density of the samples to the standard curve of theAOX level.

The AOX is a mitochondrial protein, so the mitochondria were isolated and purified from imbibed seeds following a protocol described previously (Jin *et al*., 2018). All steps involved in isolating mitochondria were conducted at 4°C. As the AOX activity is regulated by its redox state, the effects of H_2_S fumigation on the redox state of AOX were detected by reducing SDS-PAGE and Western blotting using a plant AOX antibody (AS05054 Anti-AOX, Agrisera, Sweden). By comparing the distribution of dimer AOX and monomer AOX, the changes of AOX activity were monitored.

### 2.6 Statistical analysis

Three independent biological replicates were performed on each experiment, and the results were expressed as the means ± standard error (SE). The statistically significant differences were analyzed by one-way analysis of variance (ANOVA) using SPSS (version 17, IBM SPSS, Chicago, IL, USA), and error bars were calculated based on Tukey’s multiple range test (*P* < 0.05).

## Results

### 1. Production of endogenous H_2_S was increased during seed germination

To confirm the involvement of endogenous H_2_S signaling in regulating seed germination, we assessed the time-course expression of genes responsible for endogenous H_2_S generation in seeds processing to germination, including *LCD, DES1, OASTL-A1, OASTL-B, OASTL-C, CYSC1, CYS-D1*, and *CYS-D2*. As shown in Fig. 1A, the expression of *DES1, LCD, OASTL-B*, and *OASTL-C* were up-regulated by various degrees with the extension of seed imbibition. The expression of *DES1* and *LCD* increased almost immediately after imbibition, and were maintained at higher levels in imbibed seeds. The *OASTL-C* expression showed an obvious up-regulation after 6 hours of imbibition and the *OASTL-B* expression was only slightly enhanced during germination (Fig. 1A). Notably, there were no significant changes in the expression of *CYSC1, CYS-D1*, and *CYS-D2*, genes encoding β-CAS, during germination processing (Fig. 1A). In addition, the activity of total CDes was assayed by detecting the production rate of endogenous H_2_S generated from L-cysteine and D-cysteine, and our data showed that the production rate of endogenous H_2_S raised dramatically during the first 24 hours of imbibition. Although a slight decrease occurred after 48 hours of imbibition, the H_2_S production rate was still significantly higher than that before imbibition (Fig. 1B). Consistently with the increase of the transcription of genes associated with endogenous H_2_S production and the activities of the CDes, the endogenous H_2_S content also exhibited a conspicuous increase followed by a slight decline with the process of seeds imbibition (Fig. 1B). These data suggested that imbibition stimulates the generation of endogenous H_2_S, which might act importantly in regulating seed germination.

**Fig. 1.**
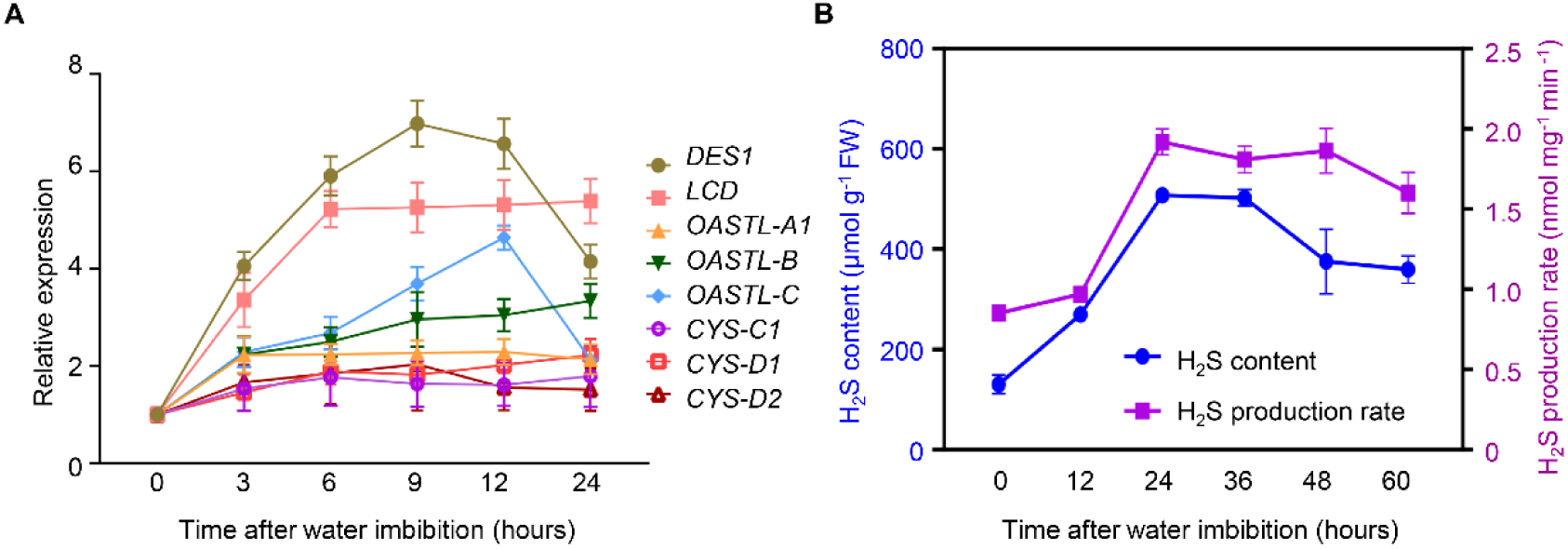
Endogenous H_2_S production was activated in imbibed Arabidopsis seeds. (A) Evolution of the expression of H_2_S generation-associated genes with the extension of imbibition; (B) The production rate and the content of endogenous H_2_S during seed germination in Arabidopsis.

### 2. H_2_S signaling participates in the regulation of seed germination

In order to investigate the significance of gasotransmitter H_2_S in regulating seed germination, we examined the effects of exogenous H_2_S at different concentrations (0, 3, 6, 9, 12, 15, 18, 21 μmol L^-1^) on the speed of seeds germination, and the sodium hydrosulfide (NaHS) was employed as exogenous H_2_S donor. As is shown in Fig. 2A, within a certain range of concentrations, exogenous H_2_S fumigation could accelerate seed germination in a concentration-dependent manner, and the positive effect of H_2_S on seed germination was strengthened with the increase of H_2_S concentration (Fig. 2A). Importantly, exogenous H_2_S fumigation with lower concentration (below 15 μmol L^-1^) can only improve the germination speed, but not the final germination rate (Fig. 2A and 2B). However, H_2_S fumigation with higher concentrations, 18 μmol L^-1^ and 21 μmol L^-1^, can obviously delay seed germination and reduce the final germination rate (Fig. 2A). Based on our data, 12 μmol L^-1^ of NaHS was used for subsequent H_2_S fumigation as exogenous H_2_S with this concentration exhibited most significant effect on promoting seed germination (Fig. 2A).

**Fig. 2.**
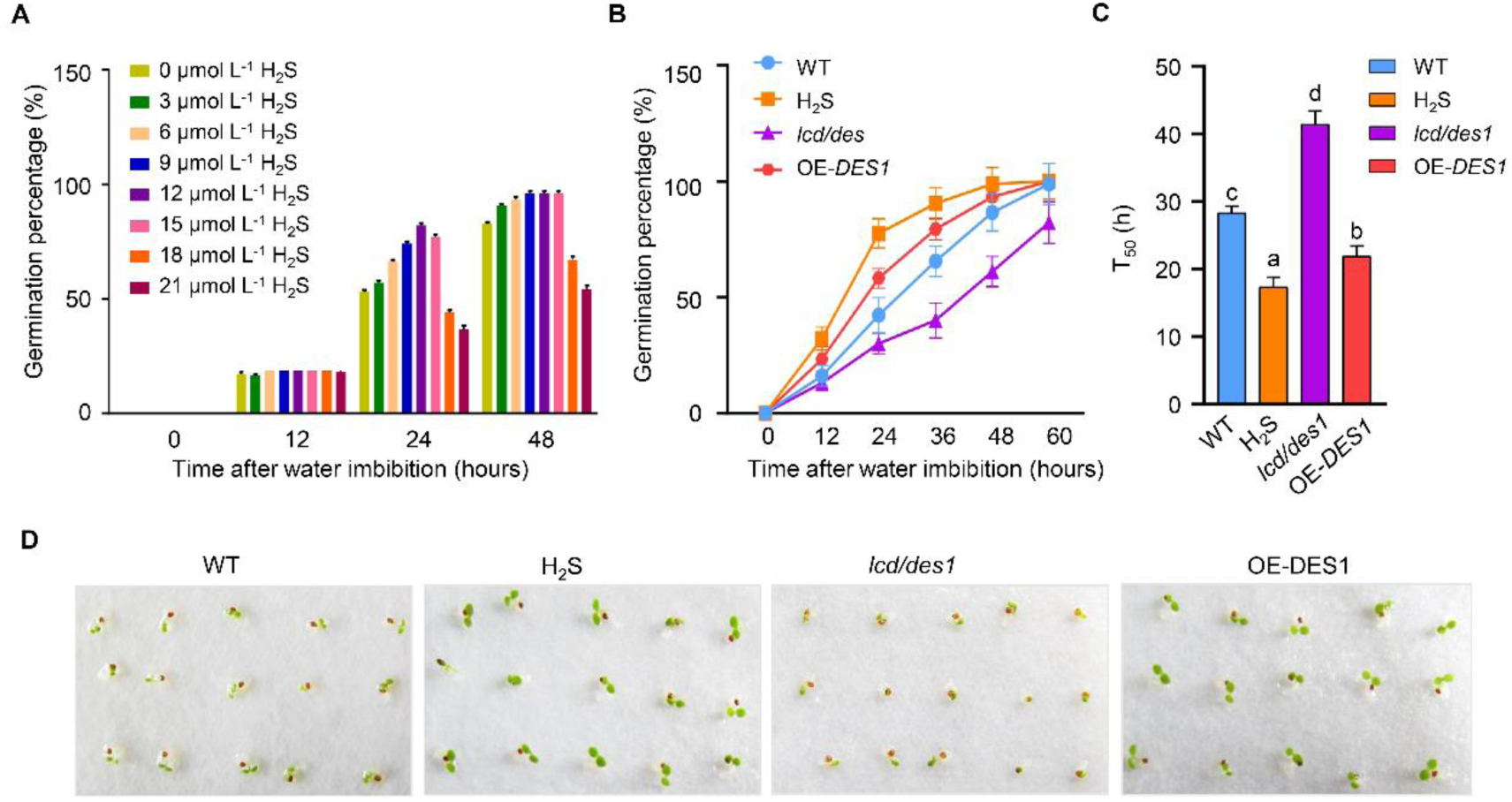
H_2_S signaling involved in accelerating seed germination in Arabidopsis. (A) Effects of exogenous H_2_S, employing the NaHS as H_2_S donor, at different concentrations (0, 3, 6, 9, 12, 15, 18, 21 μoml L^-1^) on the speed of seeds germination. (B) The germination percentages with the extension of imbibition and (C) the time to obtain 50% of germinated seeds in WT seeds, exogenous H_2_S fumigated seeds, *lcd/des1* double mutant seeds and *DES1* over-expressed (OE-*DES1*) seeds. (D) The phenotypes of germinated WT seeds, exogenous H_2_S fumigated seeds, *lcd/des1* double mutant seeds and OE-*DES1* seeds after 60 hours of imbibition.

Fumigation of WT seeds with 12 μmol L^-1^ of exogenous H_2_S could obviously accelerate seed germination by observing that H_2_S fumigation improved the percentage of germinated seeds at every time point during imbibition and significantly reduced the time to have 50% germinated seeds (T_50_) (Fig. 2B). Correspondingly, the *DES1* over-expressed transgenic seeds (OE-*DES1*) exhibited higher germination speed and faster to get 50% seeds germinated, while impairment of endogenous H_2_S accumulation by inducing mutation in *LCD* and *DES1* (*lcd/des1*) delayed germination speed and the T_50_ of *lcd/des1* seeds increased by 46.37% in comparison with WT (Fig. 2B and 2C). The phenotypes of germinated seeds after 60 hours of imbibition also confirmed the importance of H_2_S in promoting seeds germination (Fig. 2D). These data indicated that H_2_S signaling functions significantly during seed germination as exogenous H_2_S treatment or modulation of endogenous H_2_S level can obviously affect the germination speed.

### 3. Gasotransmitter H_2_S acts as a trigger of the cyanide-resistant respiration in germinating seeds

It has been reported that cyanide production would be enhanced during seed imbibition and germination (Yu *et al*., 2021). As the cyanide level affects the intensity of COX respiratory pathway through blocking electron flow, and cyanide detoxication mediated by β-CAS links the modulation of cyanide level to endogenous H_2_S generation. Therefore, the effects of exogenous and endogenous H_2_S on alternative oxidase (AOX), the respiratory terminal oxidase catalyzing the cyanide-resistant respiration, were investigated in germinating seeds. We can see from the blue column in Fig. 3 that the expression of *AOX1A, AOX1B* and *AOX1C* were up-regulated gradually with the prolongation of imbibition time (Fig. 3A, 3B, and 3C), therefore, we compared the imbibition induced activation of H_2_S signaling and AOX mediated cyanide-resistant respiration, reflected by the generation of endogenous H_2_S and the expression of *AOX* genes, respectively. As shown in Fig. 3D, the up-regulation of AOX genes expression lagged behind the increase of the expression of H_2_S-generation-associated genes and the content of endogenous H_2_S during seed germination (Fig. 3D). The expression of *AOX1A, AOX1B* and *AOX1C* were up-regulated significantly after 12 hours of imbibition (Fig. 3A, 3B, and 3C), while *DES1* and *LCD* expression as well as the endogenous H_2_S content exhibited a substantial increase almost immediately after imbibition, and followed by a longer duration of the high levels (Fig. 1A, 1B and Fig. 3D). Furthermore, exogenous H_2_S fumigation strengthened the increase of *AOX1A* and *AOX1B* expression in germinating seeds, especially in the case of *AOX1A* (Fig. 3A). Importantly, modulation of endogenous H_2_S level can also regulate the *AOX1A* expression during seed germination. The OE-*DES1* seeds showed obviously higher level of *AOX1A* transcription in comparison with WT, while *lcd/des1* seeds had decreased *AOX1A* expression in germinating seeds. These data suggested that the generation of endogenous H_2_S is induced by imbibition, and this activated H_2_S signaling functions as a trigger to up-regulate the subsequent expression of *AOX1A*, hinting that H_2_S might regulate the proportion of cyanide-resistant respiration by inducing AOX in germinating seeds.

**Fig. 3.**
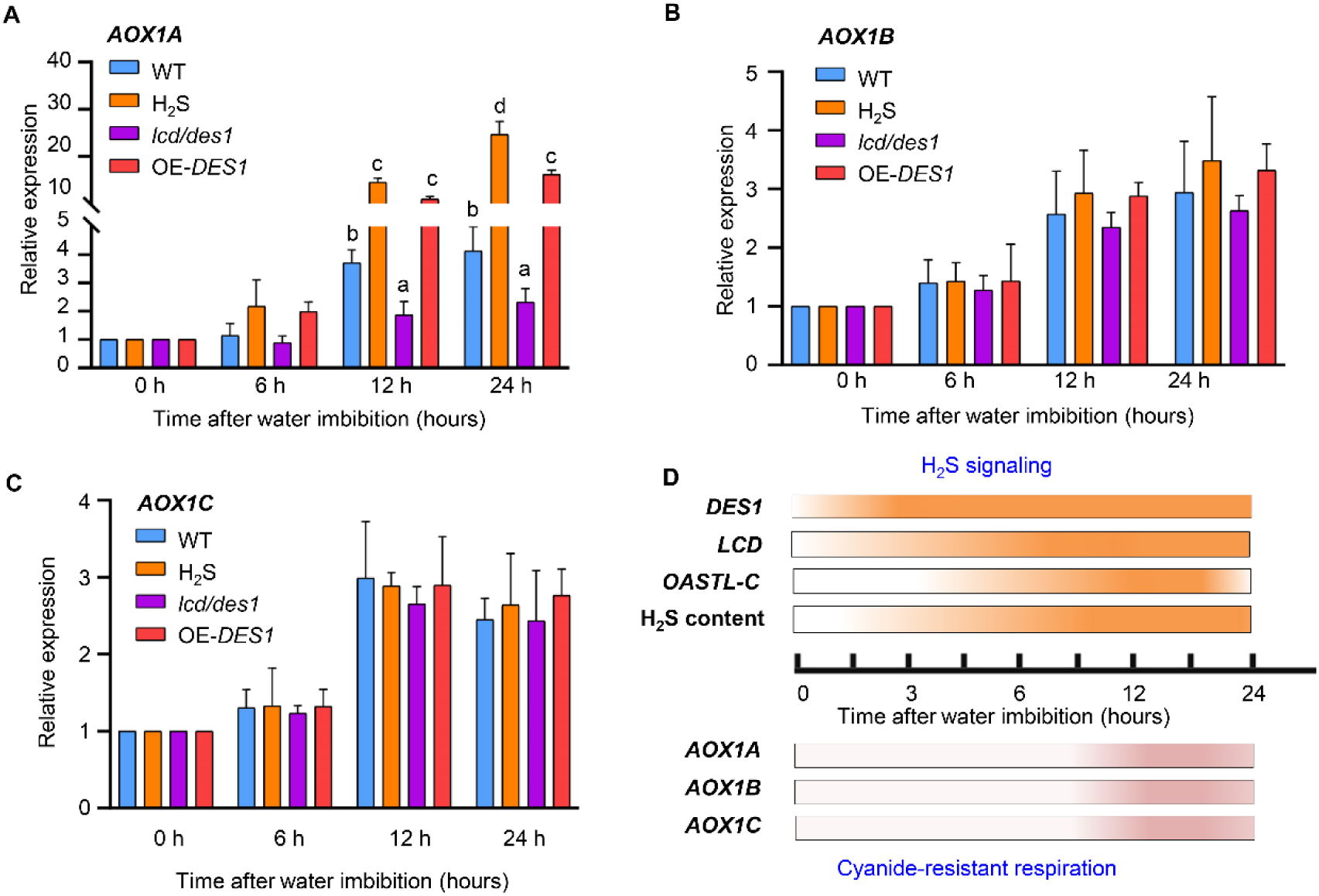
Gasotransmitter H_2_S up-regulated the transcription of AOX encoding genes. The expression of (A) *AOX1A*, (B) *AOX1B*, and (C) *AOX1C* in WT seeds, exogenous H_2_S fumigated seeds, *lcd/des1* double mutant seeds and OE-*DES1* seeds with the extension of imbibition. (D) Endogenous H_2_S signaling was activated early than the *AOX* genes expression during seed imbibition.

### 4. H_2_S regulates the cell redox state to maintain the AOX in its active and reduced state

To detect the effects of H_2_S on the protein level of AOX, we investigated the AOX concentration using the plants AOX ELISA Kit. Based on our data, it could be seen that exogenous H_2_S fumigation enhanced the abundance of AOX in germinated seeds. Correspondingly, the AOX protein level was slightly higher in the germinated OE-*DES1* seeds, and lower in the germinated *lcd/des1* seeds (Fig. 4A).

**Fig. 4.**
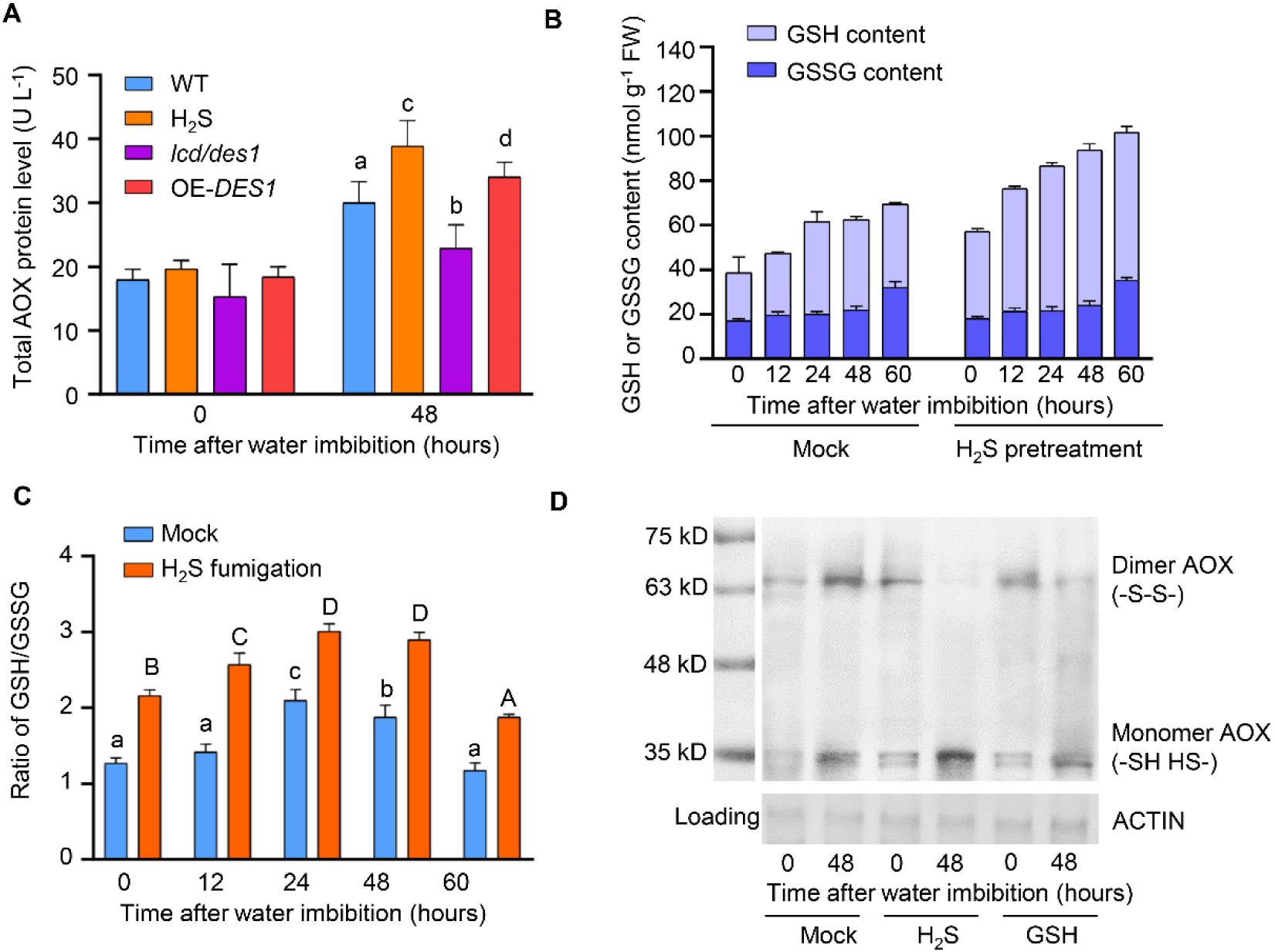
H_2_S activated the AOX activity via enhancing reduced GSH level and GSH/GSSG ratio during seed germination. (A) Effects of H_2_S signaling on the total AOX protein level in germinating seeds. (B) Effects of H_2_S fumigation on the reduced GSH and oxidized GSSG content in imbibed seeds. (B) Effects of H_2_S fumigation on the ratio of reduced GSH to oxidized GSSG in imbibed seeds. (D) Effects of H_2_S and GSH on the redox state of AOX.

It has been widely confirmed that two types of dimeric AOX structures exist. The oxidized AOX form (-S-S-) is inactivated when a highly conserved cysteine residue is oxidized to covalently link the enzyme, while the reduced AOX dimer (-SH HS-) is more active than the oxidized form (Umbach and Siedow, 1993; Moore *et al*., 1995). Therefore, the activity of AOX is greatly dependent on its redox state, correlating with the reduction of a regulatory disulfide bond (Sluse and Jarmuszkiewicz, 1998). Control of the redox state of AOX could be a powerful mechanism of regulating its activity *in vivo*, which links the activity of AOX to the general redox state of the cell (Umbach and Siedow, 1997; Sluse and Jarmuszkiewicz, 1998), so the effects of H_2_S on the redox state of AOX and the general redox state of the cell were investigated. We found that H_2_S fumigation could promote the accumulation of reduced GSH (Fig. 4B), concomitant with a marked increase of the ratio of GSH/GSSG (Fig. 4C), suggesting that gasotransmitter H_2_S might improve the cell reducing power by enhancing GSH accumulation. Moreover, the non-reducing SDS-PAGE western blot was used for detecting the dimer AOX (-S-S-) and monomer AOX (-SH HS-), and data indicated that both H_2_S and GSH treatment could enhance the level of reduced monomer AOX to various degrees, and increase the proportion of reduced AOX in germinated seeds (Fig. 4D).

Given all that, gasotransmitter H_2_S can enhance the reducing power of the cell to mediate the post-translational modification on AOX to maintain the X in its reduced and active state, which has great significance in activating the cyanide-resistant respiration during seed germination.

## Discussion

The physiological functions of H_2_S as a vital gasotransmitter in plant development and environmental response have been widely reported (Jin and Pei 2015; Corpas, 2019; Liu *et al*., 2021). Several studies have confirmed the involvement of H_2_S signaling in seed germination (Zhang *et al*., 2008; Zhang *et al*., 2010; Zhang *et al*., 2010; Li *et al*., 2012; Dooley *et al*., 2013), while the underlying mechanism of H_2_S signaling promoting seed germination remains a mystery.

In our study, fumigation with NaHS, an exogenous H_2_S donor, accelerated seed germination in a dose-dependent manner (Fig. 2A). Therefore, it was believed that within its physiological concentration scope, H_2_S can accelerate seed germination and this positive effect was reinforced with the increase of H_2_S concentration (Fig. 2A). Importantly, H_2_S treatment with physiological concentration can only improve the germination speed, but not the final germination rate (Fig. 2A and 2B). However, fumigation with high concentrations of H_2_S (18 and 21 μmol L^-1^), exceeding the physiological concentration scope, exhibited an inhibitory effect on both the germination speed and the final germination probability (Fig. 1A), so 18 and 21 μmol L^-1^ were considered as the toxic concentration of H_2_S. Endogenously, a marked up-regulation of the *DES1* and *LCD* transcription (Fig. 1A) as well as a significant enhancement of total CDes activity (Fig. 1B) were observed during seed germination, which acts importantly in affording for the increase of endogenous H_2_S content in imbibed seeds. The decelerated germination (Fig. 2B) and the increased time of obtaining 50% of germinated seeds (Fig. 2C) in *lcd/des1* seeds indicating that the H_2_S production mediated by DES1 and LCD plays a crucial role in ensuring successful germination. Collectively, our data confirmed the positive involvement of H_2_S signaling in accelerating seed germination from both the exogenous and endogenous perspectives.

β-CAS is the key enzyme for cyanide detoxification in plants, and H_2_S is accompanied by cyanide detoxification (Lai *et al*., 2009; Alvarez *et al*., 2012; Garcia *et al*., 2010). Interestingly, data in our study indicated that the expression of *CYS* genes responsible for encoding β-CAS, including *CYS-C1*, CYS-D1, and CYS-D2, exhibited a slight increase trend in germinating seeds, but the changes were not significant (Fig. 1A), suggesting that the β-CAS pathway contributes little to the increase of endogenous H_2_S production in imbibed seeds. Importantly, these data also hint the nonfeasance of cyanide detoxification during seed germination, which seems to be consistent with the functions of cyanide in seed germination reported previously. It has been reported that the emission of HCN occurs during the pre-germination period of many seeds (Esashi *et al*., 2006), and the cyanide could involve in regulating dormancy-release and seed germination (Taylorson and Hendricks, 1973; Oracz *et al*., 2008). Generally speaking, cyanide is mostly correlated with the inhibition of the terminal cytochrome oxidase (COX) in the mitochondrial respiratory pathways, so the increase of cyanide level usually implies the inhibition of COX respiratory system. Additionally, the increased endogenous H_2_S, mainly mediated by LCD and DES1, will feedback inhibit β-CAS mediated H_2_S production and might lead to the decrease of cyanide detoxification, which provides the possibility that H_2_S might regulate the proportion of cyanide-resistant respiration through modulating cyanide content. The AOX, also known as alternate oxidase, is plant-specific cyanide insensitive terminal oxidase and responsible for the activity of cyanide-resistant respiration pathway (Vanlerberghe and McIntosh, 1997; Vanlerberghe *et al*., 2010). In our study, both the gene transcription and protein abundance of AOX were enhanced in germinating seeds, suggesting that increasing of AOX respiration and partial blocking of the COX respiration might be necessary for successful seed germination.

The correlation between endogenous H_2_S production and cyanide detoxification attracts us to study the relationship between H_2_S signaling and cyanide-resistant respiration. In our study, both the endogenous H_2_S content and AOX protein level could be significantly increased by imbibition. Moreover, endogenous H_2_S generation was provoked earlier than AOX genes expression (Fig. 3A), which led us to determine the effects of H_2_S signaling on the subsequent AOX activation during seed germination. Our data verified that fumigation with 12 μmol L^-1^ NaHS extremely improved both the transcription level and protein abundance of AOX (Fig. 3B and Fig. 4 A). The *AOX1A* expression and AOX protein level were slightly enhanced in the germinating OE-*DES1* seeds while significantly weakened in the germinating *lcd/des1* seeds (Fig. 3B and Fig. 4 A). Consequently, the DES1/LCD mediated H_2_S signaling functions as a trigger to activate AOX, and thus the cyanide-resistant respiration, involving in promoting seed germination. Innovatively, our study provides important bases for the regulatory effects of H_2_S signaling on respiration pattern, which might involve improving the ratio of AOX mediated cyanide-resistant respiration, during seed germination.

It has been reported that short keto-carboxylic acids could activate AOX, among which, the pyruvate is the most effective one (Harvey *et al*., 1993; Millar *et al*., 1996). As we all known, pyruvate could be produced as a by-product in *LCD* and *DES1* mediated H_2_S production (Jin and Pei, 2015; Liu *et al*., 2021), so based on our data, the enhancement of DES1/LCD mediated H_2_S generation in imbibed seeds means that the production of pyruvate will also increase during seed germination. Therefore, it is likely that the enhanced AOX activity during seed germination could partly be attributed to pyruvate produced along with H_2_S generation. Nonetheless, the gasotransmitter H_2_S still plays a dominant role in AOX activation as the positive effect of exogenous H_2_S on activating AOX transcription and translation is significantly higher than endogenously overexpressing *DES1* (Fig. 3B and Fig. 4A).

It has been reported that H_2_S could regulate the redox state of cells by regulating both the activities of antioxidative enzymes and the contents of non-enzymatic antioxidants (Bhardwaj and Kapoor, 2021; Liu *et al*., 2021). Recently, H_2_S was found to mediate persulfidation (S-sulfidation), an oxidative post-translational modification of cysteine residues (-SH) to persulfides (-SSH), to enhance the activities of antioxidative enzymes, which is the classic enzyme-dependent pathway of ROS scavenging by H_2_S (Aroca *et al*., 2015; Aroca *et al*., 2018; Corpas *et al*., 2019; Palma *et al*., 2020; Corpas, 2019). The AOX activity is highly dependent on its redox state, so we studied the effects of H_2_S on the redox state of AOX. In our study, H_2_S could enhance the level of reduced GSH and the ratio of GSH/GSSG (Fig. 4B and 4C). Furthermore, both H_2_S and GSH could increase the content of monomer AOX (-SH HS-), and keep more AOX in its reduced and activated state (Fig. 4D). Collectively, H_2_S could act as a signal molecule to redox-regulate AOX activity during seed germination, which might correlate with the increased level of reduced GSH and the enhanced reducing power of the cell.

Based on the evidence demonstrated in this study, a novel signal model of H_2_S promoting seed germination is proposed (Fig. 5). This study presents the regulatory effects of H_2_S on respiration pattern, involving the improvement of the proportion of AOX mediated cyanide-resistant respiration, in germinating seeds. On the one hand, H_2_S could up-regulate the *AOX1A* expression to stimulate the cyanide-resistant respiration in imbibed seeds, on the other hand, H_2_S mediates the post-translational modification of AOX to keep AOX in its reduced and active state, a process that mainly involves the enhancement of the reducing power of cell by elevating the reduced GSH level and GSH/GSSG ratio. Collectively, gasotransmitter H_2_S activates AOX activity from both long-term (gene expression) and short-term (post-translational modification, allosteric activation) regulatory modes, thus enhances cyanide-resistant respiration to accelerate seed germination.

**Fig. 5.**
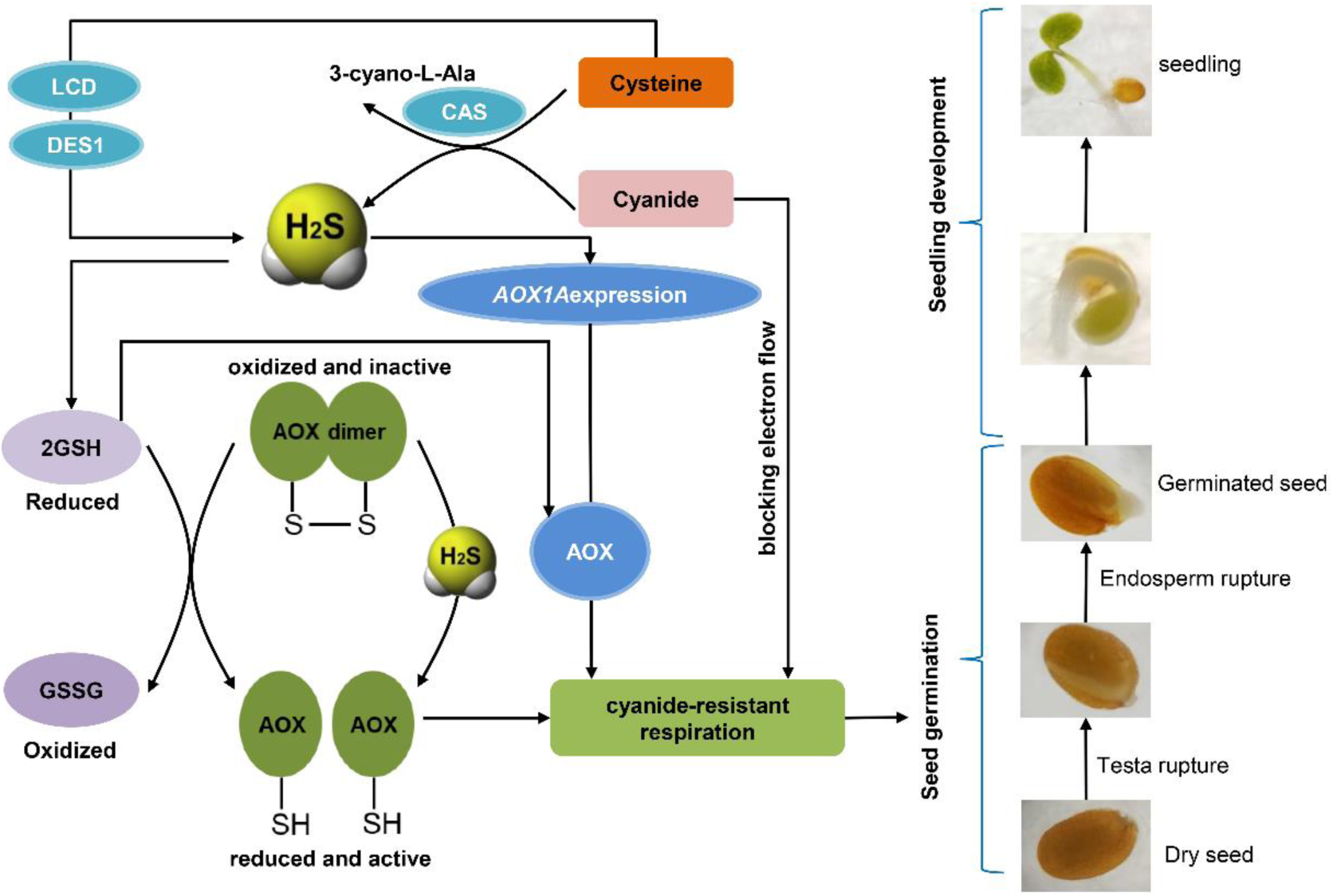
Model for gasotransmitter H_2_S promoting seed germination via enhancing AOX mediated cyanide-resistant respiration. Gasotransmitter H_2_S activates the AOX mediated cyanide-resistant respiration by up- regulating both the transcription of *AOX1A* and the protein abundance of AOX, and inducing the post-translational modification of AOX to keep more AOX in its reduced and active state through the improvement of cell reducing power and the elevation of reduced GSH level and GSH/GSSG ratio. Collectively, H_2_S activates AOX activity by both long-term (gene expression) and short-term (post-translational modification, allosteric activation) regulatory modes, thus enhances AOX mediated cyanide-resistant respiration to accelerate seed germination.

## Acknowledgments

This work is supported by a start-up fund from Zhejiang Agricultural & Forestry University (Grant No. 2020FR035), and a fund from the Natural Science Foundation of Zhejiang Province (Grant No. Q21C060002) to HF.

